# Aphid colony size is affected by plant chemotype and terpenoid mixture evenness in tansy

**DOI:** 10.1101/2025.03.04.641370

**Authors:** Annika Neuhaus-Harr, Lina Ojeda-Prieto, Xiaoyuan Zhang, Jörg-Peter Schnitzler, Wolfgang W. Weisser, Robin Heinen

## Abstract

Plants are hosts for above- and belowground insect communities that can influence each other via above-belowground plant-physiological dynamics. To mediate interactions, plants produce secondary metabolites, including terpenoids, and mixtures can differ intraspecifically. While intraspecific variation in plant chemistry gained increased interest, the extent to which intraspecific differences in plant chemistry mediate above-belowground interactions of herbivores remains unclear. We used a full factorial design with six distinct terpenoid chemotypes, differing in their chemical diversity of tansy (*Tanacetum vulgare*). We exposed these to the aboveground herbivore *Macrosiphoniella tanacetaria* (Hemiptera: Aphididae), the belowground herbivore *Agriotes* sp. (Coleoptera: Elateridae), no herbivore or both herbivores, to determine if chemotypes or the chemical diversity of plant compounds affected aphid performance and if the chemical profile mediated the interactions between herbivores. We found that aphid colony size differed between chemotypes, with the strongest colony increase over time in a mixed-mixtures chemotype, and the weakest in a β-thujone chemotype. Root herbivory had no effect on aphid colony size and this did not differ between chemotypes. Aphid colony size was positively correlated with terpenoid evenness, but not with other diversity components. Tansy chemotypes differed in their morphological responses to aboveground herbivory, whereas belowground herbivory exerted minimal impacts. Overall, our results show that intraspecific variation in terpenoid profiles directly and indirectly modify ecological interactions on a plant, with plant chemistry mediating aphid performance and chemotypes differing in their morphological responses to herbivory.

## Introduction

Plants play a central role in multi-layered interactions, serving as hosts for complex insect communities across trophic levels. Specialised plant metabolites are important for regulating mediating interactions between plants and their living environment (Agrawal & Weber, 2015). Within a single plant species, individuals can exhibit differences in specialised metabolite profiles (Weng et al., 2021), and this intraspecific variation can lead to significant differences in the outcome of interactions within plant species (Bączek et al., 2019; Christensen et al., 2019; Kleine & Müller, 2011; Rahimova, Neuhaus-Harr, et al., 2024; Schoonhoven et al., 2005). How different aspects of plant chemical profiles, particularly their diversity, relate to ecological plant interactions is currently receiving a lot of interest (Jakobs & Müller, 2018; Kessler & Kalske, 2018; Petrén, Anaia, et al., 2023; Petrén, Köllner, et al., 2023; Richards et al., 2015; Wetzel & Whitehead, 2020; Whitehead et al., 2021; Ziaja & Müller, 2023). For instance, in a recent study, the diversity and distinct composition of terpenoid mixtures in tansy plants (*Tanacetum vulgare*) affected host preference of specialised tansy aphids in choice assays (Neuhaus-Harr et al., 2024). Aphids preferred the chemotypes dominanted by α-thujone/β-thujone and β-trans-chrysanthenyl acetate, while avoiding the chemotype with a mixed terpenoid profile (Neuhaus-Harr et al., 2024).

Chemical diversity can be described in a number of ways, including by the distinct difference of chemical profiles, but can further be described by its three main diversity components: richness, evenness, and disparity (Petrén, Köllner, et al., 2023). Chemical richness, a straightforward measure of phytochemical diversity, refers to the number of compounds in a tissue. It is hypothesised that chemically richer plants benefit when having e.g. multiple herbivore species as attackers, compared with plants that produce fewer compounds (Junker, 2016). Chemical evenness describes the number of compounds and takes into account their relative abundance. Evidence also exists for chemical evenness to affect interactions between plants and insects. For example, specialised tansy aphids tend to avoid tansy plants with higher terpenoid evenness levels (Neuhaus-Harr et al., 2024). Chemical disparity considers the qualitative differences of tissues in terms of chemical compounds that are present, but to date very few studies have taken the ecological role of chemical disparity into account (Petrén, Köllner, et al., 2023). Though numerous studies provide valuable insights into different aspects of plant chemistry and its role in ecology (Dyer, 2018; Junker, 2018), we still lack a comprehensive understanding of how different components of plant chemical diversity shape plant-insect interactions and which aspects are most relevant as mediators of plant-herbivore interactions (Petrén, Köllner, et al., 2023).

While the effects of secondary metabolites on plant-herbivore interactions are documented, less is known about how intraspecific differences in chemical profiles affect the interactions between multiple simultaneous attackers on the same plant, especially if these herbivores feed on different plant parts. It is plausible that plant chemotype composition may determine the outcome of above-belowground herbivore interactions on the same plant. Aboveground and belowground herbivores can induce local and/or systemic defences in plants, leading to altered plant metabolism, changes in plant morphology, or resource allocation towards defence (Lehndal & Ågren, 2015; Maron & Crone, 2006; Zhou et al., 2015). This, in turn, can affect herbivores feeding on other plant parts. For example, root-feeding herbivores such as the endo-parasitic nematode *Pratylenchus penetrans* or the larvae of the cabbage root fly *Delia radicum* significantly alter the nutritional quality of plant shoots in *Brassica nigra*, through changes in glucosinolate levels, which in turn negatively affect the growth and reproduction rate of caterpillars of the small cabbage white, *Pieris rapae* (Van Dam et al., 2005). According to a meta-analysis, the outcome of above-belowground herbivore interactions depends on multiple factors such as herbivore feeding guild (Johnson et al., 2012). For instance, belowground chewing larvae of beetle species had a positive effect on aboveground Homoptera, such as aphids, but a negative effect on aboveground Hymenoptera (Johnson et al., 2012). Furthermore, Yang and colleagues (2024) recently suggested that species-specific plant responses to herbivores are more important than herbivore identity or herbivore specialization in determining the plant response to sequential attacks. How these interactions are affected by intraspecific differences in plant chemistry, are not yet fully understood.

Insect herbivores, above- and below-ground typically have multiple negative effects on plants. In their review, Nabity and colleagues (2009) point out that herbivory reduces photosynthetic rates due to tissue loss and disruption of photosynthesis around the missing tissue. Herbivory also reduces plant size, growth, and seed production (Hodkinson & Hughes, 1982; Myers & Sarfraz, 2017). It remains unclear, how chemical diversity of plants might mitigate these effects, as plants that differ in their chemical composition could also differ in their resistance and resilience to above- or belowground herbivory.

This study uses *Tanacetum vulgare* L. (Asteraceae), a perennial plant known for its variable aromatic terpenoid composition. Tansy has a wide geographical distribution and hosts a diverse community of herbivores, including aphids with varying host specificity (Keskitalo et al., 2001; Kleine & Müller, 2011; Schmitz, 1998). Tansy plants are characterised by their richness in mono- and sesquiterpenoids and can be classified into chemotypes based on their terpenoid composition (Keskitalo et al., 2001; Kleine & Müller, 2011). It is hypothesised that specialised aphids have adapted to the potentially harmful metabolites in tansy and may even use plant volatiles to locate their hosts (Jakobs & Müller, 2019; Schoonhoven et al., 2005). Aphid preference, colonisation, growth rate, survival, and genotype structure have been partially attributed to the chemotypes of tansy (Benedek et al., 2015; Clancy et al., 2018; Neuhaus-Harr et al., 2024; Senft et al., 2017, 2019; Zytynska et al., 2019). For example, it has been found that when given the choice between different chemotypes, the tansy aphid *Macrosiphoniella tanacetaria* preferred the two chemotypes dominated by trans-chrysanthenyl acetate (Chrys_acet) and α-thujone/β-thujone (Athu_Bthu) over the others (Neuhaus-Harr et al., 2024).

Using six biologically replicated *Tanacetum vulgare* chemotypes that differ in their leaf terpenoid composition, total terpenoid concentration, terpenoid richness, terpenoid evenness and Shannon diversity, we test the effects of the presence of generalist belowground root herbivores (wireworm larvae: a mixture of *Agriotes lineatus* and *Agriotes obscurus*) on the aboveground herbivore performance of the tansy aphid *Macrosiphoniella tanacteria* and whether chemotypes mitigate these relationships. Furthermore, we test whether the effects of herbivory on the plant morphology differ between chemotypes. We address the following hypotheses:

**(H1)** We expect aphids to perform best on the chemotypes they preferred in choice assays in a previous study (Neuhaus-Harr et al., 2024).

**(H2)** Belowground coleopteran herbivores will positively affect aphid colony size and colony growth, but these relationships will differ in their strength and direction between chemotypes.

**(H3)** More chemically diverse plants (i.e., higher total terpenoid concentration, higher terpenoid richness, higher terpenoid evenness, and higher terpenoid Shannon diversity index) will result in smaller aphid colonies but the interaction with belowground treatment will modify this relationship.

**(H4)** Above- and belowground herbivores will have a detrimental effect on plant growth and morphology, but the strength of these effects differs across chemotypes. Specifically, plants infested with both herbivores will have the least chlorophyll content in their leaves, grow less tall, and have lower dry weight compared to plants with only one or no herbivore, but we predict that chemically less diverse plants will suffer less from herbivory as they possibly use more resources towards growth and not defence.

## Methods & Materials

### Plant Material

In 2019, tansy plants were collected in Jena, Germany, and their terpenoid profiles were analysed to determine chemotypes (described in Neuhaus-Harr et al. 2024). Briefly, leaf material was freeze-dried, homogenised and weighed and by adding one-bromodecane as internal standard, terpenoids were extracted in heptane. Extracts were centrifuged and by using gas chromatography and mass spectrometry, supernatants were analysed with Helium as carrier gas, using an alkane standard mix as a reference. Retention indices, and mass spectra were compared with compounds in Pherobase (El-Sayed, 2012), entries of the National Institute of Standards and Technology 2014 and mass spectra reported in Adams (2017). Using unsupervised hierarchical k-means clustering with the ‘hclust()’ function, the plants were grouped into seven clusters (k = 7). Further details regarding the characterization of these established chemotype lines are described in Neuhaus-Harr et al. 2024. Chemotypes varied in their dominant compound(s), total terpenoid concentration, terpenoid richness, terpenoid evenness, and terpenoid Shannon diversity, as presented in the supplementary information and detailed in a previous study (Table S4, Neuhaus-Harr et al. 2024). Chemotype terpenoid profiles ranged between 21 – 29 terpenoid compounds, and based on their relative concentration, diversity components were calculated for each daughter plant from their terpenoid profiles, using the ‘vegan’ package (Oksanen et al., 2022).

### Propagation of plant material

In May 2022, 40 shoot cuttings were taken from the specific chemotypes tansy plants, which are maintained in a common garden in Freising, Germany. The stems of fresh plants were cut into parts with 1-2 cm below and 4-5 cm above a leaf node. The leaf size was reduced by clipping the pinnate leaves to decrease evaporation and the risk of mould. The cuttings were then planted into seedling trays filled with standard potting substrate (Stender potting substrate C 700 coarse structure, 1 kg NPK minerals m–3, pH 5.5–6.0). All cuttings were kept in a greenhouse with bottom watering and additional lighting (16:00:8:00 h L:D) following standard protocols described in Neuhaus-Harr 2024. Three weeks later, 25 rooted cuttings from each chemotype were transplanted into individual 11 cm-diameter pots. To maintain a target electrical conductivity of 1.0, the plants were fertilised with Universol Blue fertiliser (18% N – 11% P – 18% K; ICL Deutschland). In July, all plants were repotted into 19 cm pots to avoid pot limitation. Clones from the same daughter line were grown in pots randomly distributed over different tables in the greenhouse to avoid initial growth bias due to environmental variation within the greenhouse. After the plants were well established, we placed the pots into a covered vegetation hall with iron mesh (5 cm) walls. From each chemotype, we randomly selected 40 established cuttings (with 13-14 cloned individuals from each of the three chemotype-specific daughter lines; see Table S5). Clones were obtained by taking stem cuttings from the respective daughter, which were propagated as mentioned above.

### Experimental design

We established a fully factorial design with either no herbivore, only the aboveground herbivore (aphid *Macrosiphoniella tanacetaria*), only the belowground herbivore (wireworm *Agriotes* sp), or both herbivores. Each plant was *a priori* assigned to one of four different treatments and arranged in a block design with 10 replicated blocks, totalling 240 plants (2 aboveground treatment levels (aphid/no aphid) x 2 belowground treatment levels (wireworm/no wireworm) x 6 chemotypes x 3 biologically replicated daughters each with 3 or 4 clonal replicates; 10 total; see Table S5).

For the belowground herbivory treatment, wireworms (a mixture of *Agriotes lineatus* and *Agriotes obscurus*) were obtained in 2022 from Wageningen University, Lelystad, The Netherlands. Upon arrival, the wireworms were kept in sandy soil at 20°C with two sliced potatoes as a food source until they were used in the experiment. For the aboveground herbivory treatment, we collected *M. tanacetaria* aphids from Jena, Germany. Aphids were kept in cages in a climate-controlled lab at room temperature with supplemental light (16:00: 8:00 h L:D) provided by two tubes (T5 FQ 80W/865 HO High Output LUMILUX Daylight G5, OSRAM GmbH, Munich, Germany) and with 2-4 tansy chemotype plants obtained from local Freising populations. A minimum of 100 adult aphids were collected and transferred to Petri dishes with fresh leaves to generate age-specific cohorts. One day later, all adults were removed, and the remaining aphid nymphs were kept in the dishes for three more days in a Fitotron standard growth chamber (21/16°C, 60% RH, Weiss Technik). Aphid cohorts were supplied with fresh leaves daily until they were used in the experiment.

### Above- and belowground treatments

Four days before aphid infestation (day 0; Fig. 1), two 1 cm deep holes were made in the soil surface in all plant pots, and those pots assigned to the belowground treatment were infested with two wireworms each. During the experiment, all plants were placed on a plant saucer to prevent wireworms from escaping. After the belowground treatment was started, all plants were watered twice daily with up to 400 ml water per watering event, depending on the plant’s water demand and soil humidity.

**Fig. 1.**
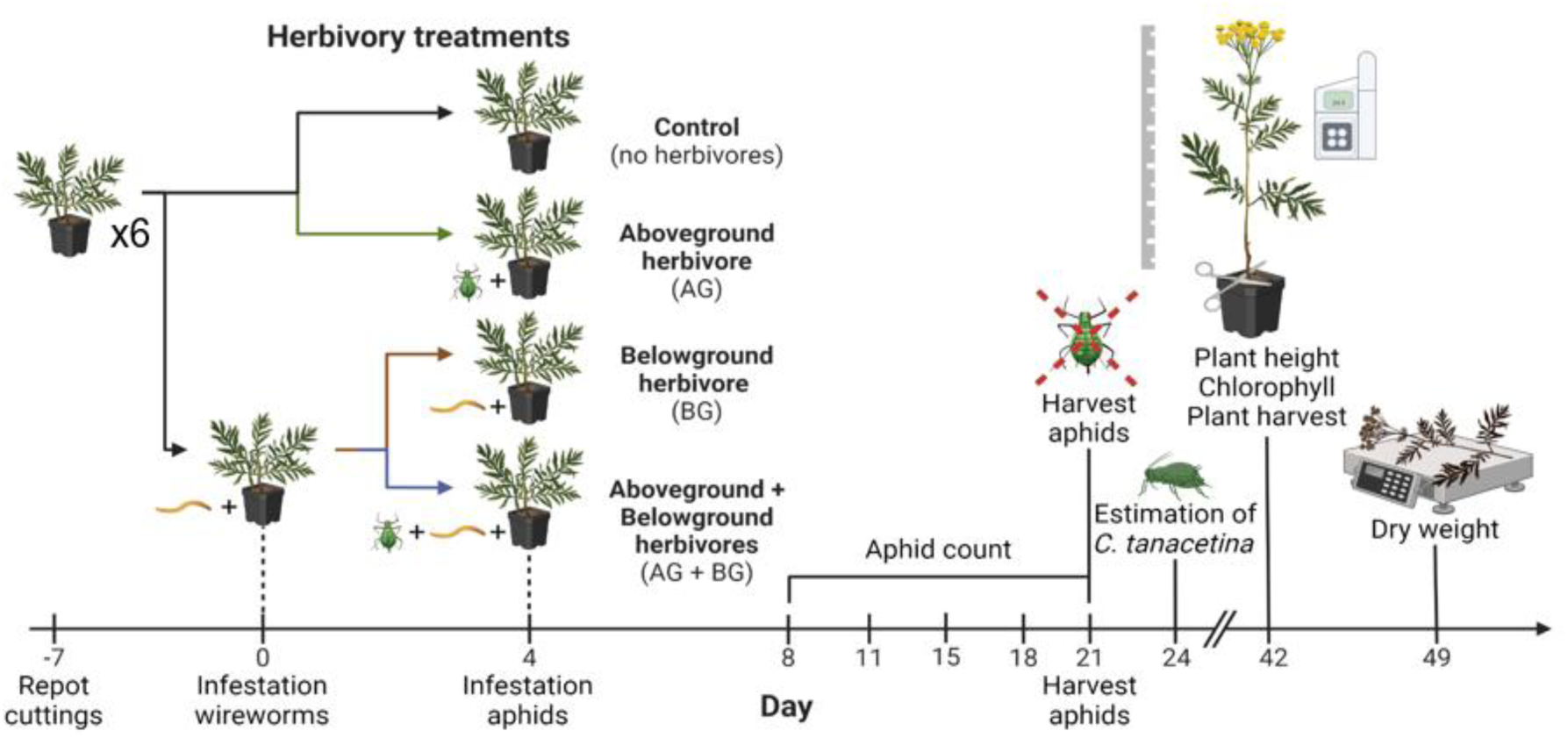
Experimental design and timeline of the above-belowground herbivore experiment. The experimental timeline, including the last repotting event, infestation with wireworms and aphids, aphid counts, assessment of infestation levels with *Coloradoa tanacetina* and measurements of plant height, chlorophyll and dry weight. Created with Biorender®.

On the day of the aphid treatment (Fig. 1; day 4), we carefully attached a fine mesh bag (11 cm x 9.5 cm) with breathable and see-through fabric on the second youngest, fully expanded leave, without squeezing the leaf or petiole. We did this to every plant, to maintain consistency in leaf and bag placement and reduce potential differences in mesh bag effects. Plants that were allocated to the aphid treatment additionally received two three-day-old aphid nymphs inside their mesh bag. The mesh bags protected the nymphs from predators and kept the colony in place, allowing for controlled observations.

### Insect measurements

The experimental timeline started on the day during which plants with belowground treatment were infested with two wireworms (day 0, Fig. 1), which occurred one week after the last repotting event. Aboveground treatment (infestation with two three-day *M. tanacetaria* in a closed mesh bag on the second youngest fully expanded leaf) was performed on day 4. The colony size of *M. tanacetaria* was counted on days 8, 11, 15, 18, and 21. On day 21, *M. tanacetaria* aphids were harvested from each plant. The numbers of *Coloradoa tanacetina* aphids were estimated on day 24.

After harvesting the plants (day 49, Fig. 1), we traced back wireworms by going through the soil by hand, and noted that a disproportionate number had pupated or even enclosed as adults – which is not common in such belowground treatments (R. Heinen, pers. obs.), as wireworms typically spend several years in larval stage (Furlan, 1998). This high level of pupation may have been caused by a series of heatwaves that took place during the experiment. For this reason, at the end of the experiment, we counted the number of larvae, pupae, and adult beetles from each pot to ensure that no wireworms were missing and we could account for it statistically. We retrieved 68 wireworms that had pupated or even reached adulthood, while 90 wireworms remained in their larval stadium. Seventy-four individuals went missing and could not be found back, which may indicate that they reached adulthood and escaped the pots. We tested the effect of these pupation events on treatment efficacy in separate models (referred to below as Model A for belowground treatment and Model B for retrieved number of larvae).

### Plant measurements

Three days after harvesting the leaves with infested aphids, we noted whether plants were infested by the tansy leaf margin aphid *Coloradoa tanacetina* and estimated infestation numbers to statistically assess the effect of these unplanned infestations on *M. tanacetaria* performance and plant morphology. *C. tanacetina* estimation was done by picking three random leaves per plant and calculating the average number of aphids in steps of 10 for numbers between 0 and 100, in steps of 50 for numbers above 100, up to 300. On day 42 (Fig. 1), once plant growth stagnated, we assessed plant height by measuring the distance from the soil to the highest point of the plant, without straightening the plant. We further measured the average chlorophyll content of three random leaves per plant using a chlorophyll meter (Konica Minolta SPAD-502Plus, Tokyo, Japan) as a proxy of plant health. Aboveground dry weight was assessed after drying the samples for 78 hours at 60°C. Furthermore, we calculated total terpenoid concentration (unit), terpenoid richness, terpenoid evenness, and terpenoid Shannon diversity from the absolute terpenoid profiles for each daughter using the ‘vegan’ package (Oksanen et al., 2022).

### Statistical analysis

All statistical analyses were performed in R version 4.1.2 (R core Team 2021). We used linear mixed models, as detailed below, to test our hypotheses with the ‘lmer()’ command from the ‘lme4’ package (Bates et al., 2014). As aphid counts were strongly left-skewed, we square root-transformed this variable in every model to meet model assumptions. The assumptions of all models were assessed by plotting QQ plots, residual plots, and scale-location plots. We used ‘Anova()’ from the ‘car’ package to calculate p-values (Fox & Weisberg, 2019). All model codes can be found in the supplement Table S6.

To address **H1**, whether chemotypes would drive aphid colony size, we created a model where we included the final aphid count as the response variable and chemotype as a fixed factor. To test whether natural colonisation of the experimental plants by the aphid *C. tanacetina* affected *M. tanacetaria* colony size, we included *C. tanacetina* abundance as a covariate in this model. Block and daughter ID were included as random effects to account for variation between blocks and clonal replicates.

To simultaneously test **H1** and **H2**, we created two model variants to test the effect of belowground treatment (Model variant A), or the number of retrieved larvae (Model variant B) on aphid colony size over time. Model variant A included chemotype, belowground treatment, observation day, and their interactions as fixed effects. Block, daughter ID to account for variation between blocks and clonal replicates. Furthermore, unique plant ID, nested in observation day, was used as a random effect, to account for the fact that aphids were counted more than once on the same plant over time. In Model variant B we replaced belowground treatment with the number of retrieved wireworm larvae to investigate whether pupation during the experimental procedure affected the treatment effect on aphid colony size.

To address **H3**, whether components of plant chemical diversity (terpenoid richness, terpenoid Shannon diversity, terpenoid evenness, total terpenoid concentration) mediate the effect of wireworms on aphid colony size, we set up two multiple regression models. In Model variant A, we included belowground treatment and all chemical diversity components as fixed effects. Block was included as a random effect. In Model variant B, we replaced belowground treatment with the number of retrieved wireworm larvae to investigate whether pupation during the experimental procedure affected the treatment effect on aphid colony size. In the next step we used variance inflation factors, ‘vif()’ to account for multicollinearity in our models. In both models, we excluded terpenoid Shannon diversity, as it highly correlated with terpenoid evenness.

To address **H4** whether above- or belowground herbivory affected plant morphology (i.e., chlorophyll, plant height, plant biomass) we used linear mixed models. In Model variant A, we included *C. tanacetina* as a covariate, treatment (aboveground herbivory, belowground herbivory, above- and belowground herbivory, and control), chemotype, and the interaction of these three variables as fixed factors. Block and daughter ID were included as random effects. As the retrieved number of larvae were not homogeneously distributed across all treatment and chemotype combinations, this limited our analytical power at this level for a Model B variant. Therefore, we tested the effect of retrieved wireworm larvae to investigate whether pupation during the experimental procedure affected the treatment effect on plant variables. Block and daughter were included as random effects.

## Results

### Experimental procedure

We infested 120 plants with aphids, two of which died during the experiment and were excluded from analysis. On 37 out of the remaining 118 plants, no aphids survived until the end of the aphid assay. In three of these plants, predatory mirid bugs were found in the mesh bags that were installed to protect the aphids. Hence, we excluded these three plants from further analyses. After three weeks, the remaining aphid colonies ranged from 1 to 77. We tested whether aphid survival differed between chemotypes. We found that aphid survival did not significantly differ between chemotypes (χ^2^_5_ = 7.26, p >0.05; see Table S7). Within chemotypes the survival rates between daughters varied significantly (χ^2^_1_ = 11.30, p < 0.001, see Table S7). As aphid colonies that consist of zero individuals throughout the experiment provide no meaningful insights in aphid colony growth on different chemotypes, we analysed aphid colony growth data with and without the non-surviving aphids included. As patterns did not differ between the two approaches, we present the aphid colony growth data in the main text excluding the non-surviving aphids. However, for transparency we also present all analyses including the non-surviving aphids in the supplementary information (Table S8, S9, Fig. S5, S6).

Over the course of the experiment *Coloradoa tanacetina* aphids naturally occurred on our experimental plants, with colony sizes ranging from zero (on 25 plants) up to 200 individuals (on 3 plants) per leaf, and tested for each variable, whether its presence had a significant effect. Where appropriate, *C. tanacetina* numbers were included as covariates.

### Aphid performance across chemotypes (H1) and belowground treatment (H2)

We found no evidence that the degree of naturally occurring infestation by *C. tanacetina* affected *M. tanacetaria* final aphid colony size, neither when belowground treatment was included (χ^2^_1_ = 2.09, p > 0.05; see Table S10 Model A) nor when the number of wireworms were included instead of the belowground treatment (χ^2^_3_ = 1.53, p > 0.05; see Table S10 Model B). Furthermore, the numbers of *C. tanacetina* did not differ across chemotypes (χ^2^_5_ = 1.49, p > 0.05; see Table S11), colony sizes of *M. tanacetaria* (χ^2^_1_ = 2.21, p > 0.05; see Table S11) or the interplay of chemotype and *M. tanacetaria* colony sizes (χ^2^_1_ = 7.66, df = 5, p > 0.05; see Table S11). For this reason, *C. tanacetina* numbers were not further included in the further aphid analyses below. A graph with *C. tanacetina* abundance for all chemotypes can be found in the supplement (Fig. S7).

Four days after adding two aphids to plants with aphid treatments, we counted the aphid numbers. After eleven days, aphids had matured and first offspring was recorded. On day 24, the colony sizes ranged from one to 77. Colony size of *M. tanacetaria* significantly increased with time the experiment (Model A, day: χ^2^_1_ = 183.72, p < 0.001; Table 1). Aphid colony size also significantly differed between chemotypes over time, indicated by the interaction between day and chemotype (Model A, chemotype * day: χ^2^_5_ = 17.92, p = 0.003; Table 1), and visible as different slopes in Fig. 2a). Aphid colony sizes increased faster on chemotypes with a mixed terpenoid profile, particularly the mixed chemotype with low terpenoid concentration. In line with this, final aphid colony sizes were higher on the mixed low chemotype than on the others (Fig. 2b). However, belowground herbivory treatment did not affect aphid colony size (Table 1). When the number of retrieved wireworm larvae was included in the model instead of belowground herbivory treatment, we observed very similar patterns (Model B, day: χ^2^_1_ = 168.97, p < 0.001; chemotype * day: χ^2^_5_ = 13.43, p = 0.020; Table 1). Note that plants with zero aphids were excluded. A table with zero aphids included can be found in the supplementary (Table S8).

**Fig. 2.**
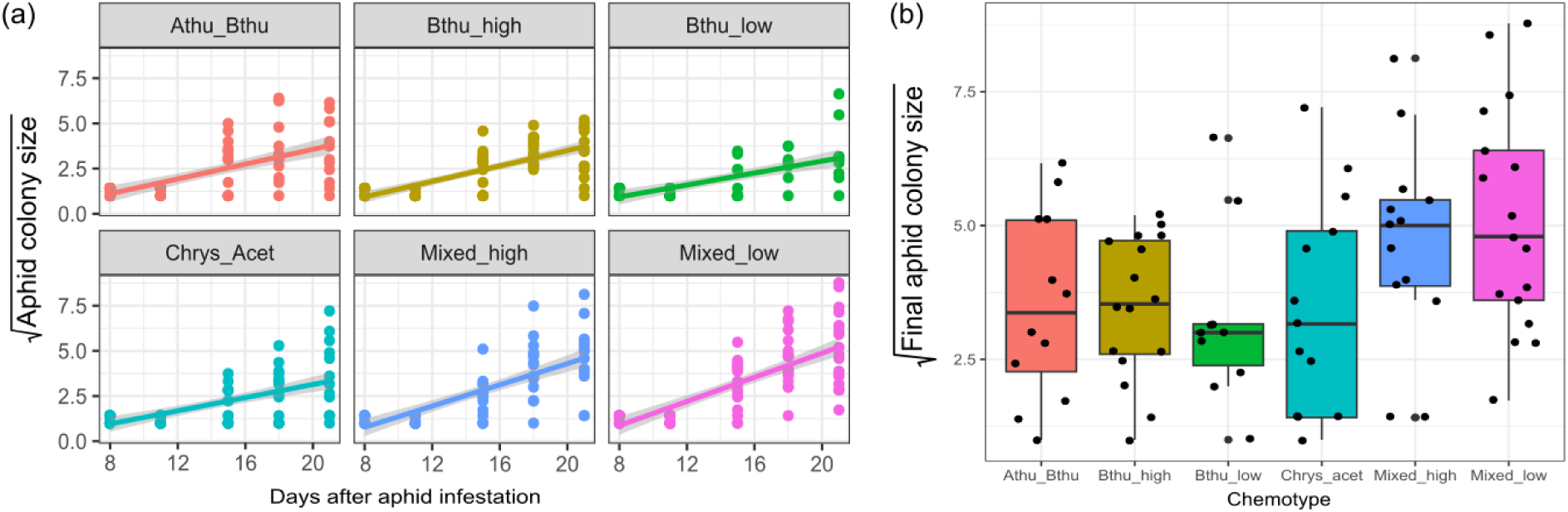
(a) Square root-transformed *M. tanacetaria* colony size over time in days after aphid infestation, across chemotypes. (b) Final aphid colony size at the time of the experimental harvest for different tansy chemotypes. Boxes represent the variation in data, where the lower hinge corresponds to the first quartile (25th percentile) and the upper hinge depicts the third quartile (75th percentile). Whiskers indicate the 5% and 95% percentiles; solid lines within boxes represent the medians. Black dots indicate individual sample values. The six chemotypes are depicted in different colours for convenience. Note that in both graphs plants with zero aphids were excluded. Graphs with zero aphids included can be found in the supplementary (Fig. S5)

**Table 1:**
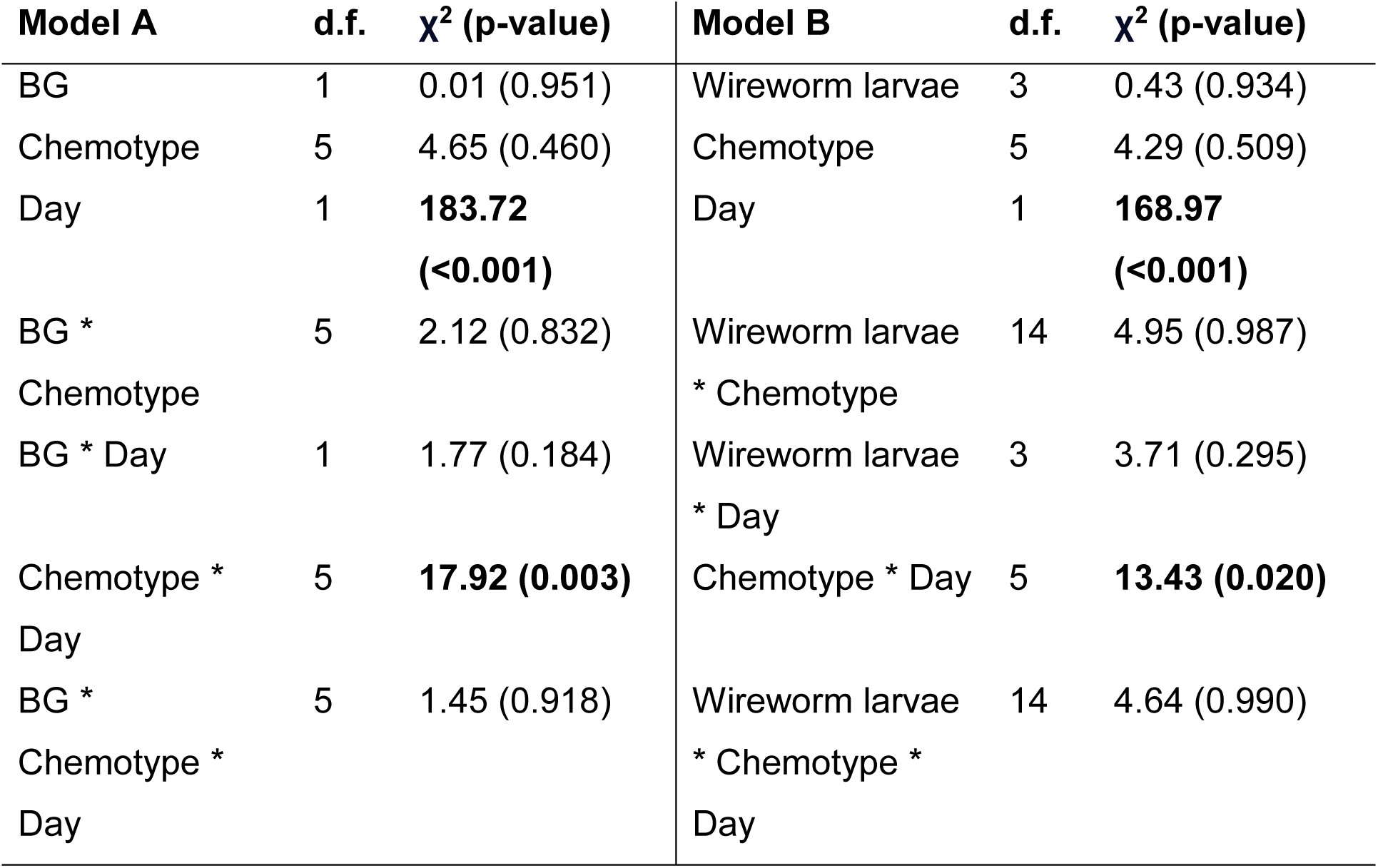
Output from a mixed linear model for *M. tanacetaria* colony size over time, using either belowground herbivory treatment (BG; Model A) or the number of retrieved wireworm larvae (Wireworm larvae; Model B), and the day and chemotype, and the interaction terms as fixed effects. In both models, block, daughter ID, and individual ID (nested within day) were used as random effects.

### Chemical diversity components (H3) and their effects on aphid colony size

When investigating the relationships between different components of chemical diversity, we found that *M. tanacetaria* colonies were significantly larger on plants that had a higher leaf terpenoid evenness (χ^2^_1_ = 4.34, p = 0.037; Table 2; Fig. 3a). We observed no effects for terpenoid richness, or total terpenoid concentration (Table 2). Terpenoid Shannon diversity was excluded as this is highly correlated to terpenoid richness and evenness. There was no effect of the belowground herbivory treatment on aphid colony size.

**Fig. 3.**
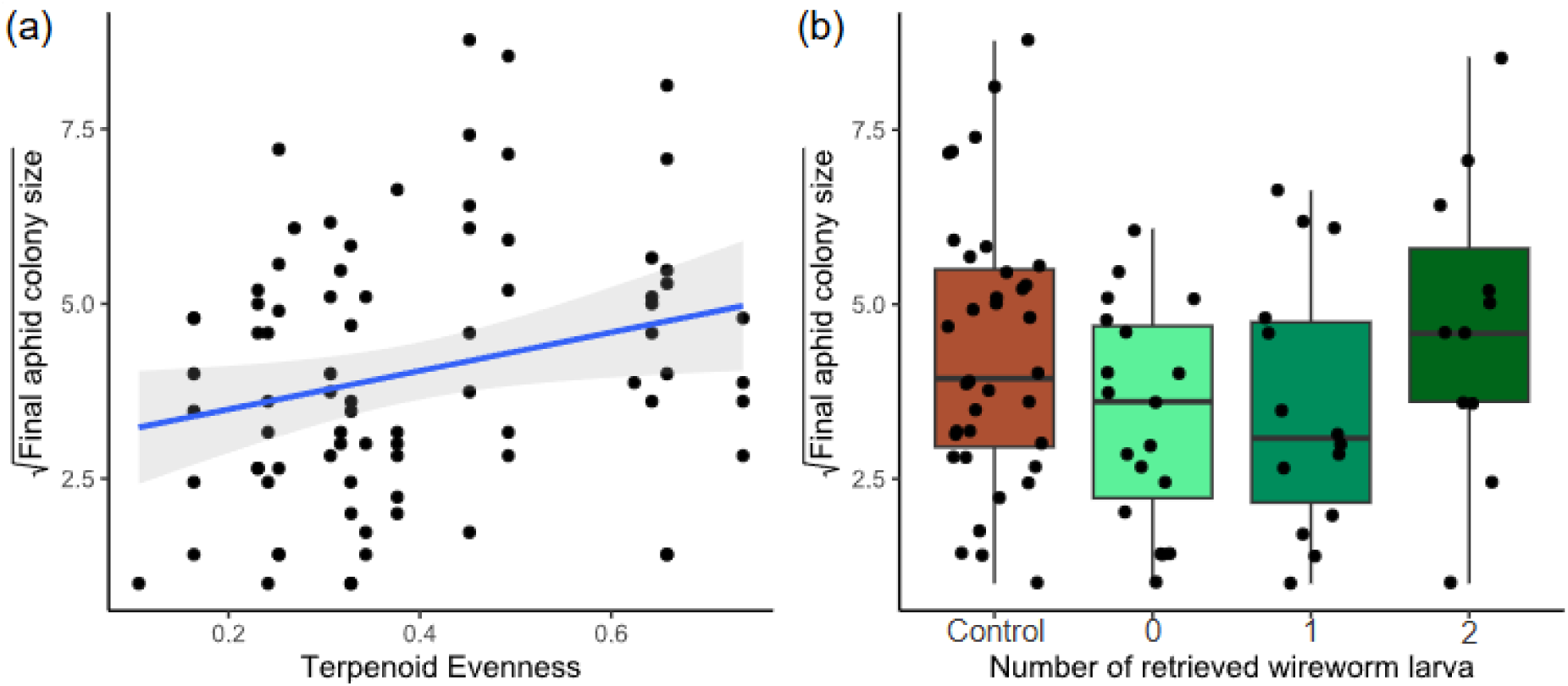
(a) Square root-transformed *M. tanacetaria* colony size on plants differing in leaf terpenoid evenness. The linear trendline depicts average predicted values based on a linear model, and the shaded area depicts the 95% confidence interval. (b) Box plots visualizing square root-transformed *M. tanacetaria* colony size on plants with no added wireworms, compared to plants on which 0, 1 or 2 wireworm larvae were retrieved after the harvest. Boxes represent the variation in data, where the lower hinge corresponds to the first quartile (25th percentile) and the upper hinge depicts the third quartile (75th percentile). Whiskers indicate the 5% and 95% percentiles; solid lines within boxes represent the medians. Black dots indicate individual sample values. Note that in both graphs plants with zero aphids were excluded. Graphs with zero aphids included can be found in the supplementary (Fig. S6)

**Table 2:** Output from a mixed linear model for final *M. tanacetaria* colony size, using belowground herbivory treatment (BG; Model A) or the number of retrieved wireworm larvae (Wireworm larvae; Model B), and terpenoid richness, terpenoid evenness and total terpenoid concentration calculated based on the terpenoid profile of the 18 daughter plants (three for each of the six chemotypes) as fixed effects and the block as random effect.

| Model A | d.f. | $\chi^2$ (p-value) | Model B | d.f. | $\chi^2$ (p-value) |
| --- | --- | --- | --- | --- | --- |
| BG | 1 | 2.22 (0.136) | Wireworm larvae | 3 | 4.77 (0.189) |
| Evenness | 1 | **4.34 (0.037)** | Evenness | 1 | 3.21 (0.073) |
| Richness | 1 | 0.04 (0.837) | Richness | 1 | 0.09 (0.764) |
| Concentration | 1 | 1.97 (0.160) | Concentration | 1 | 1.60 (0.206) |

When including the number of retrieved wireworm larvae instead of belowground herbivory treatment (Model B), we found that terpenoid evenness was nearly significant (χ^2^_1_ = 3.21, p > 0.05; Table 2). Wireworm larvae did not affect final aphid colony sizes, regardless if we checked for treatments (Table 2 Model A) or included the number of wireworm larvae retrieved (Table 2, Model B, Fig. 3b). Note that plants with zero aphids were excluded. A table with zero aphids included can be found in the supplementary (Table S9).

### Effect of above- and belowground herbivory treatments on plant morphology (H4)

We observed that the strength of infestation by *C. tanacetina* did not affect aboveground plant dry weight, but significantly negatively affected plant height and marginally affected chlorophyll content of the experimental plants, and hence was kept as a covariate in all plant response models below (Table 3; Fig. S8).

**Table 3:** Output from a mixed linear model for average leaf chlorophyll content [units], plant height [cm], and aboveground dry weight [gr] taking the *C. tanacetina* abundance, treatment (Aboveground herbivory, Belowground herbivory, both herbivory treatments, and control) and chemotype as fixed effects and block and daughter ID as random effects.

|  | Plant dry weight |  | Plant height |  | Chlorophyll content |  |
| --- | --- | --- | --- | --- | --- | --- |
| | d.f. | $\chi^2$ (p-value) | d.f. | $\chi^2$ (p-value) | d.f. | $\chi^2$ (p-value) |
| *C. tanacetina*<br>abundance | 1 | 0.22 (0.638) | 1 | **13.42 (<0.001)** | 1 | 2.90 (0.089) |
| AG | 1 | 0.50 (0.480) | 1 | **7.75 (0.005)** | 1 | 0.82 (0.364) |
| BG | 1 | 0.33 (0.563) | 1 | 0.33 (0.569) | 1 | 1.54 (0.215) |
| Chemotype | 5 | 1.36 (0.929) | 5 | 2.26 (0.812) | 5 | **18.77 (0.002)** |
| AG * BG | 1 | 2.42 (0.12) | 1 | 1.03 (0.311) | 1 | **6.30 (0.012)** |
| AG * Chemotype | 5 | **11.70 (0.039)** | 5 | **13.59 (0.018)** | 5 | 6.99 (0.230) |
| BG * Chemotype | 5 | 6.10 (0.296) | 5 | 2.84 (0.725) | 5 | 9.16 (0.103) |
| AG * BG *<br>Chemotype | 5 | 2.49 (0.778) | 5 | 3.42 (0.636) | 5 | 6.75 (0.240) |

We found that the interaction of aboveground treatment and chemotype significantly affected the aboveground plant dry weight (χ^2^_5_ = 11.70, p = 0.039; Table 3). Specifically, plants from the Mixed_high chemotype had a higher aboveground dry weight when they received an aboveground treatment with aphids, compared to control plants, while in all other chemotypes aboveground treatment with aphids had either a negative or no effect on plant aboveground dry weight (Fig. 4a). Plant height was significantly affected by aboveground treatment (χ^2^_1_ = 7.75, p = 0.005; Table 3), and its interaction with chemotype (X^2^_5_ = 13.59, p = 0.018; Table 3). While control plants grew taller for most chemotypes than those exposed to aboveground treatment with aphids, plants from the Mixed_low chemotype grew taller in the aboveground treatment, compared to control plants (Fig. 4b). The average leaf chlorophyll content significantly differed across chemotypes (χ^2^_5_ = 18.77, p = 0.002, Table 3) and was also affected by an interaction between above- and below-ground treatments (χ^2^_1_ = 6.30, p = 0.012; Table 3). Specifically, plants exposed to belowground treatment with wireworms seemed to have a higher chlorophyll content than plants that received aboveground treatment with aphids, or than control plants (Fig. 4c). Further, the Bthu_low chemotype had a significantly lower average leaf chlorophyll content than all other chemotypes, whereas the Mixed_low chemotype showed the highest average leaf chlorophyll content (Fig. 4d).

**Fig. 4.**
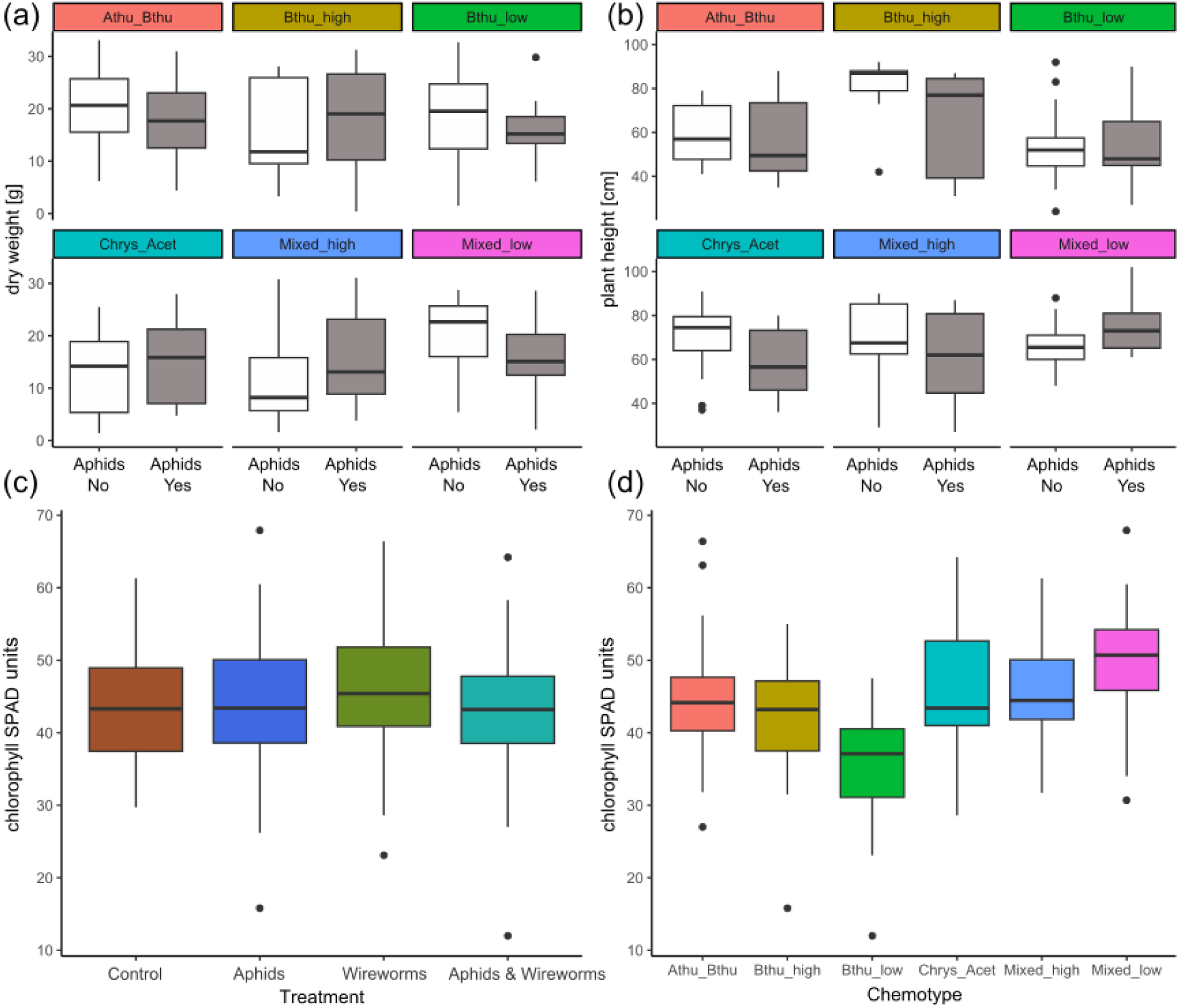
Effects of plant chemotype, aboveground (aphid) and belowground (wireworm) treatment on a) plant dry weight, b) plant height, c,d) chlorophyll content. White boxes represent plants without aphids, grey boxes represent plants with aphids. Panel (c) represents an interactive effect between above- and belowground treatment on average leaf chlorophyll content (SPAD units), and (d) depicts differences in chlorophyll content across chemotypes. Boxes represent the variation in data, where the lower hinge corresponds to the first quartile (25th percentile) and the upper hinge depicts the third quartile (75th percentile). Whiskers indicate the 5% and 95% percentiles; solid lines within boxes represent the medians. Black dots indicate outliers

## Discussion

In this study, we tested the effects of chemotypic variation in leaf terpenoid profiles on the interactions between a belowground root herbivore and an aboveground phloem feeding aphid in tansy (*T. vulgare*). We found that aphid colony size development over time significantly differed across chemotypes. Contrary to our expectations, belowground infestation with wireworms did not have any effect on aphid colony size. Therefore, chemotypes did not mediate interactions between belowground herbivores and aboveground herbivores. Our multiple regression models indicated a positive relationship between tansy leaf terpenoid evenness and final aphid colony size, suggesting that aphids perform better when the compounds in the terpenoid mixtures are more evenly distributed in concentration. We found that aphid presence significantly affected plant dry weight and plant height, but that the patterns differed between chemotypes. Root herbivore presence had surprisingly little effect on plant growth of any chemotype. Taken together, our results suggest a strong role of plant chemotype as a determinant of aphid colony dynamics, likely driven by the distribution of the relative abundance of terpenoid compounds in the mixtures.

In line with our first hypothesis, we found that *Macrosiphoniella tanacetaria* colony size development significantly differed between different *T. vulgare* chemotypes. This is in line with a previous study showing that this aphid species was significantly affected by an interaction between tansy chemotype and plant part (Jakobs & Müller, 2018), but in our study, the effects of chemotypes were substantially more pronounced than in the aforementioned. Interestingly, a previous study that used the same tansy chemotype lines as used in the present study found that, when given a choice, *M. tanacetaria* adults preferred to feed on leaves from the chemotype Athu_Bthu compared to Bthu_high or Mixed_low, and generally showed a higher attraction towards the chemotypes Athu_Bthu and Chrys_acet (Neuhaus-Harr et al., 2024). This is interesting, because this indicates that what aphids prefer to feed on does not seem to match how they perform on it. According to the “mother knows best” or “preference-performance” hypothesis, adult insects should prefer plants on which their offspring have maximum performance, which is believed to be true for many aboveground specialist insects (Birke & Aluja, 2018; Gripenberg et al., 2010). This raises important questions regarding the costs and benefits of attraction to specific chemistry in aphids: Why would aphids choose low-quality hosts over high-quality hosts, and how do chemical cues relate to plant qualitative components?

The previous prediction that belowground herbivores should positively influence aboveground herbivores (Masters et al., 1993), has since been challenged in many subsequent studies indicating that above- and belowground interactions are highly context dependent (Johnson et al., 2012). In line with this, but contrary to our second hypothesis, we did not find that belowground herbivores had a positive influence on *M. tanacetaria* colony size. There could be several explanations for this. First, although Coleoptera (such as wireworm larvae) as belowground herbivores typically have a positive influence on aboveground Homoptera (e.g., aphids) (Johnson & Murray, 2008), this effect was only found when the both herbivores arrived at the same time, which indicates that the plant response is early, and potentially short-lived (Erb et al., 2011; Johnson et al., 2012). As we infested plants with wireworms three days before aphids, plants might have already recovered from the root attack and the increase of leaf nutrients due to herbivore stress (which benefits the aphids), had already faded out (Johnson et al., 2012). Second, when ending the experiment and retrieving the wireworms, we found that many of them had pupated over the course of the experimental duration, or, in some cases, had even turned into adults. This is possibly due to local heat waves that occurred during the experiment, in August 2022. Although wireworms typically live for many years, the warm conditions may have sped up their larval cycle, as temperature is typically negatively correlated to the length of larval life cycles in insects (Furlan, 1996; Meikle & Patt, 2011). As wireworms do not feed during pupation, the resulting levels of herbivory might have been too low to have a significant effect on the aboveground aphids. It could be that wireworm feeding on tansy roots is not consistent, although we have observed in unpublished pilots that wireworms readily feed on plant roots, and particularly on the fine root hairs (J-P Schnitzler, pers. obs.). This calls for further studies with higher levels of root-herbivores. A final explanation may be that responses to root herbivory in tansy are local, rather than systemic. As we did not find differences among chemotypes, this could indicate that belowground and aboveground plant responses might be compartmentalised. As described in a recent study, tansy terpenoid profiles differ strongly between above- and below-ground compartments, following different biosynthetic pathways (Rahimova, Heinen, et al., 2024). It is possible that there is minimal resource allocation or defence pathways overlap. Further studies unravelling how and where wireworm feeding affects plant physiological processes is needed to draw definitive conclusions.

We predicted that more chemically diverse plants (i.e., higher total terpenoid concentration, higher terpenoid richness, higher terpenoid evenness, and higher terpenoid Shannon diversity) would be more strongly defended, which we hypothesized would lead to reduced *M. tanacetaria* colony size. Further, we predicted that belowground treatment with root herbivores would modify this relationship. Contrary to our predictions, we found that *M. tanacetaria* colonies were significantly larger on plants with higher terpenoid evenness. No significant effects were observed for the other chemodiversity components. There was no effect of belowground herbivory treatment, nor any interaction between belowground treatment and the chemical diversity components on aphid colony size. The role of evenness in ecological contexts is highly dependent on the organisms and functions involved (Petrén, Köllner, et al., 2023). If ecological functions rely on compounds occurring in similar abundances, a high evenness could lead to stronger synergistic interactions among these compounds (Petrén, Köllner, et al., 2023). Such synergies could benefit aphid growth, which might explain why higher terpenoid evenness correlated with larger aphid colonies in our study. Conversely, if specific functions, like suppressing aphid growth, depend on a few key compounds, high evenness could dilute the relative abundance of these critical compounds, resulting in more favourable conditions for aphids.

Although several studies show positive relationships between chemical profiles dominated by individual compounds, these results seem to suggest that across a range of different chemotypes, single compound-dominated mixtures (i.e., low evenness) are detrimental to aphid colony development. One important caveat is that evenness may also be partly confounded by disparity, i.e., the effect of the origin of the compound on its ecological effect (Petrén, Anaia, et al., 2023). For instance, in our study, low-evenness profiles were typically dominated by β-thujone, or by chrysanthenyl acetate, which, although both monoterpenoids derived from geranyl diphosphate, are the result of different downstream pathways, and may have different ecological effects on herbivores (Rahimova, Neuhaus-Harr, et al., 2024). Similarly, other compound dominated mixtures in nature may have even different impacts on aphids. Disentangling the effects of evenness from the effect of disparity would require large-scale sampling and propagation efforts to test the effects of the full natural breadth of terpenoid mixtures and their diversity on aphid colonies under standardised conditions, and this would be an important direction for the future.

We found that the infestation of tansy by *M. tanacetaria* influenced plant height and plant dry weight, although the direction of the effect differed between chemotypes. The leaf chlorophyll content also differed between chemotypes and was lower when a plant experienced above- and belowground herbivory. This implies that chemotypes might differ in their growth and defence strategies as has been found in multiple other plant species (He et al., 2022; Huot et al., 2014; Züst & Agrawal, 2017). Both of the mixed chemotypes, i.e., the chemotypes with the highest richness, diversity and evenness of compounds, grew either taller or had a higher dry weight when infested with aphids, compared to control plants. Interestingly, also these chemotypes had the largest *M. tanacetaria* colony size. Although it may seem counterintuitive, perhaps plants with more diverse terpenoid chemotypes may save resources by the production of a diverse mixtures of compounds in low relative abundance, allowing them to invest resources into growing and compensating for loss through herbivory, while other chemotypes invest more into chemically defending themselves through production of dominant compounds. As producing chemical defence is costly, it is often associated with a restriction in growth (Havko et al., 2016; Herms & Mattson, 1992; Huot et al., 2014; Sestari & Campos, 2022), which has been found in many plant species (Campos et al., 2016; Haak et al., 2012; Hayashi et al., 2020; Mihaliak & Lincoln, 1989). However, recent research shows that growth vs. defense is not simply a consequence of limited resources but a strategy of plants to maximise their fitness, that is context-dependent and aims to ensure greatest fitness of a plant in its environment (Campos et al., 2016; Guo et al., 2018; Kliebenstein, 2016). As we found differences between chemotypes, this implies that individuals within species might display very different strategies and that this might be closely connected to secondary metabolites. A cost-benefit analysis of the maintenance of chemical diversity, for instance relative to other well-characterized processes in plants, would greatly help us understand chemical profiles in the context of defence optimization strategies.

To conclude, we found that intraspecific plant chemistry plays an important role in how plants interact with their biotic and abiotic environment. Secondary metabolites not only serve as a defence system, through repelling herbivores or attracting herbivore predators, but also seem to be connected to other life history traits such as plant growth. This study might help us understand the role of chemotypes in the growth-defence trade-off of aboveground herbivory. While belowground herbivory had a small effect on the plant and none on the aboveground herbivore, these effects did not differ between plants with different leaf chemotypes. This may be an indication that plant defence is locally compartmentalised, as the chemotypic profile of roots highly differs from that found in leaves (Rahimova, Heinen, et al., 2024). It might be that minimal above-ground defence signalling takes place in this system for this reason. Our study sheds light on the role of plant chemotypes on mitigating above- and belowground herbivory, but we call for further research on root and shoot chemistry and their respective roles in governing above-belowground insect-plant interactions.

## Acknowledgements

We thank the German research foundation DFG for funding this study. This work was funded through grants of the project 415496540 to WWW (WE3081/40-1), as part of the Research Unit (RU) FOR 3000 and by the project 245400135 to WWW (WE3081/25-2). We also thank the staff at the Technical University of Munich (TUM) Plant Technology Center Dürnast, particularly Robert Hansel, Sabine Zuber and Petra Scheuerer, for providing excellent care during plant propagation and preparation phase.

## Statements & Declarations

### Author Contributions

All authors contributed to the study conception and design. Annika Neuhaus-Harr, Lina Ojeda-Prieto and Robin Heinen originally formulated the idea, with input from Xiaoyuan Zhang, Wolfgang W. Weisser and Jörg-Peter Schnitzler. Xiaoyuan Zhang, Annika Neuhaus-Harr and Lina Ojeda-Prieto conducted fieldwork with help from Robin Heinen. Annika Neuhaus-Harr led the formal analyses with substantial input from Lina Ojeda-Prieto and Robin Heinen. Annika Neuhaus-Harr wrote the manuscript with substantial input from Robin Heinen and Lina Ojeda-Prieto. Xiaoyuan Zhang, Wolfgang W. Weisser and Jörg-Peter Schnitzler provided editorial advice.

