## Supplemental Tables 4-11, Supplemental figures 5-8 for "Aphid colony size is affected by plant chemotype and terpenoid mixture evenness in tansy"

Supplementary information

**Table S4:** Overview of all 18 daughters (three daughters per chemotype), their dominant compound(s), their terpenoid shannon diversity, terpenoid evenness, terpenoid richness and total terpenoid concentration.

| Chemotype | Daughter | Dominant compound | Shannon diversity | Even-ness | Rich-ness | Total conc. |
|---|---|---|---|---|---|---|
| Bthu_low | 101 | β-thujone | 0.33 | 0.11 | 23 | 30.94 |
| Bthu_low | 103 | β-thujone | 1.15 | 0.38 | 21 | 36.61 |
| Bthu_low | 104 | β-thujone | 1.07 | 0.32 | 29 | 45.09 |
| Mixed_high | 21 | Mixed (borneol, cis-sabinene hydrate, eugenol, sabinene, others) | 1.93 | 0.62 | 22 | 51.48 |
| Mixed_high | 23 | Mixed (borneol, cis-sabinene hydrate, eugenol, sabinene, others) | 2.15 | 0.66 | 26 | 55.08 |
| Mixed_high | 30 | Mixed (borneol, cis-sabinene hydrate, eugenol, sabinene, others) | 2.03 | 0.64 | 23 | 57.60 |
| Chrys_Acet | 94 | Chrysantenyl acetate | 0.91 | 0.27 | 29 | 58.66 |
| Chrys_Acet | 95 | Chrysantenyl acetate | 0.80 | 0.24 | 28 | 54.52 |
| Chrys_Acet | 100 | Chrysantenyl acetate | 0.82 | 0.25 | 26 | 55.64 |
| Athu_Bthu | 55 | α-thujone; β-thujone | 1.03 | 0.33 | 23 | 59.94 |
| Athu_Bthu | 56 | α-thujone; β-thujone | 1.01 | 0.31 | 27 | 51.61 |
| Athu_Bthu | 57 | α-thujone; β-thujone | 1.13 | 0.34 | 27 | 49.90 |
| Bthu_high | 11 | β-thujone | 1.06 | 0.33 | 25 | 52.37 |
| Bthu_high | 14 | β-thujone | 0.51 | 0.16 | 22 | 44.44 |
| Bthu_high | 17 | β-thujone | 0.72 | 0.23 | 23 | 52.14 |
| Mixed_low | 64 | Mixed (sabinene, α- | 1.60 | 0.49 | 26 | 24.71 |

| | | | | | | |
|---|---|---|---|---|---|---|
| | | pinene, others) | | | | |
| Mixed_low | 67 | Mixed (sabinene, α-pinene, others) | 2.40 | 0.74 | 26 | 30.87 |
| Mixed_low | 68 | Mixed (sabinene, α-pinene, others) | 1.42 | 0.45 | 23 | 23.91 |

**Table S5**: Overview of all 240 plants that were used for the experiment. Three daughters per each of the six chemotypes were used. Ten replicates per chemotype were assigned to each treatment and randomly distributed over ten blocks.

| Chemotype | Daughter | ID | Aphids | Wireworms (WW) | Treatment | Block |
|---|---|---|---|---|---|---|
| Bthu_low | 101 | 16 | Yes | No | Aphids | 9 |
| Bthu_low | 101 | 4 | Yes | No | Aphids | 8 |
| Bthu_low | 101 | 9 | Yes | No | Aphids | 2 |
| Bthu_low | 101 | 12 | Yes | Yes | Aphids + WW | 8 |
| Bthu_low | 101 | 18 | Yes | Yes | Aphids + WW | 2 |
| Bthu_low | 101 | 7 | Yes | Yes | Aphids + WW | 9 |
| Bthu_low | 101 | 10 | No | No | Control | 2 |
| Bthu_low | 101 | 14 | No | No | Control | 9 |
| Bthu_low | 101 | 8 | No | No | Control | 8 |
| Bthu_low | 101 | 11 | No | Yes | WW | 10 |
| Bthu_low | 101 | 17 | No | Yes | WW | 8 |
| Bthu_low | 101 | 3 | No | Yes | WW | 9 |
| Bthu_low | 101 | 5 | No | Yes | WW | 2 |
| Bthu_low | 103 | 11 | Yes | No | Aphids | 3 |
| Bthu_low | 103 | 3 | Yes | No | Aphids | 5 |
| Bthu_low | 103 | 4 | Yes | No | Aphids | 10 |
| Bthu_low | 103 | 9 | Yes | No | Aphids | 1 |
| Bthu_low | 103 | 1 | Yes | Yes | Aphids + WW | 3 |
| Bthu_low | 103 | 10 | Yes | Yes | Aphids + WW | 5 |
| Bthu_low | 103 | 14 | Yes | Yes | Aphids + WW | 1 |
| Bthu_low | 103 | 6 | Yes | Yes | Aphids + WW | 10 |
| Bthu_low | 103 | 15 | No | No | Control | 5 |
| Bthu_low | 103 | 2 | No | No | Control | 10 |
| Bthu_low | 103 | 5 | No | No | Control | 3 |

| | | | | | | |
|---|---|---|---|---|---|---|
| Bthu_low | 103 | 8 | No | No | Control | 1 |
| Bthu_low | 103 | 12 | No | Yes | WW | 5 |
| Bthu_low | 103 | 13 | No | Yes | WW | 3 |
| Bthu_low | 103 | 7 | No | Yes | WW | 1 |
| Bthu_low | 104 | 15 | Yes | No | Aphids | 4 |
| Bthu_low | 104 | 3 | Yes | No | Aphids | 6 |
| Bthu_low | 104 | 9 | Yes | No | Aphids | 7 |
| Bthu_low | 104 | 10 | Yes | Yes | Aphids + WW | 4 |
| Bthu_low | 104 | 5 | Yes | Yes | Aphids + WW | 6 |
| Bthu_low | 104 | 7 | Yes | Yes | Aphids + WW | 7 |
| Bthu_low | 104 | 2 | No | No | Control | 6 |
| Bthu_low | 104 | 20 | No | No | Control | 4 |
| Bthu_low | 104 | 6 | No | No | Control | 7 |
| Bthu_low | 104 | 1 | No | Yes | WW | 7 |
| Bthu_low | 104 | 12 | No | Yes | WW | 6 |
| Bthu_low | 104 | 16 | No | Yes | WW | 4 |
| Mixed_high | 21 | 10 | Yes | No | Aphids | 10 |
| Mixed_high | 21 | 13 | Yes | No | Aphids | 7 |
| Mixed_high | 21 | 5 | Yes | No | Aphids | 9 |
| Mixed_high | 21 | 1 | Yes | Yes | Aphids + WW | 9 |
| Mixed_high | 21 | 11 | Yes | Yes | Aphids + WW | 10 |
| Mixed_high | 21 | 2 | No | No | Control | 7 |
| Mixed_high | 21 | 6 | No | No | Control | 9 |
| Mixed_high | 21 | 7 | No | No | Control | 10 |
| Mixed_high | 21 | 17 | No | Yes | WW | 10 |
| Mixed_high | 21 | 3 | No | Yes | WW | 9 |
| Mixed_high | 21 | 4 | No | Yes | WW | 7 |
| Mixed_high | 23 | 17 | Yes | No | Aphids | 1 |
| Mixed_high | 23 | 20 | Yes | No | Aphids | 5 |
| Mixed_high | 23 | 5 | Yes | No | Aphids | 2 |
| Mixed_high | 23 | 9 | Yes | No | Aphids | 3 |
| Mixed_high | 23 | 1 | Yes | Yes | Aphids + WW | 3 |
| Mixed_high | 23 | 19 | Yes | Yes | Aphids + WW | 5 |
| Mixed_high | 23 | 2 | Yes | Yes | Aphids + WW | 1 |
| Mixed_high | 23 | 6 | Yes | Yes | Aphids + WW | 2 |
| Mixed_high | 23 | 10 | No | No | Control | 1 |
| Mixed_high | 23 | 11 | No | No | Control | 3 |

| | | | | | | |
|---|---|---|---|---|---|---|
| Mixed_high | 23 | 12 | No | No | Control | 5 |
| Mixed_high | 23 | 15 | No | No | Control | 2 |
| Mixed_high | 23 | 14 | No | Yes | WW | 5 |
| Mixed_high | 23 | 16 | No | Yes | WW | 3 |
| Mixed_high | 23 | 18 | No | Yes | WW | 1 |
| Mixed_high | 23 | 8 | No | Yes | WW | 2 |
| Mixed_high | 30 | 10 | Yes | No | Aphids | 8 |
| Mixed_high | 30 | 17 | Yes | No | Aphids | 6 |
| Mixed_high | 30 | 19 | Yes | No | Aphids | 4 |
| Mixed_high | 30 | 13 | Yes | Yes | Aphids + WW | 7 |
| Mixed_high | 30 | 16 | Yes | Yes | Aphids + WW | 8 |
| Mixed_high | 30 | 20 | Yes | Yes | Aphids + WW | 6 |
| Mixed_high | 30 | 8 | Yes | Yes | Aphids + WW | 4 |
| Mixed_high | 30 | 11 | No | No | Control | 8 |
| Mixed_high | 30 | 12 | No | No | Control | 4 |
| Mixed_high | 30 | 15 | No | No | Control | 6 |
| Mixed_high | 30 | 14 | No | Yes | WW | 8 |
| Mixed_high | 30 | 18 | No | Yes | WW | 6 |
| Mixed_high | 30 | 2 | No | Yes | WW | 4 |
| Chrys_Acet | 94 | 11 | Yes | No | Aphids | 1 |
| Chrys_Acet | 94 | 15 | Yes | No | Aphids | 2 |
| Chrys_Acet | 94 | 2 | Yes | No | Aphids | 9 |
| Chrys_Acet | 94 | 1 | Yes | Yes | Aphids + WW | 9 |
| Chrys_Acet | 94 | 18 | Yes | Yes | Aphids + WW | 2 |
| Chrys_Acet | 94 | 5 | Yes | Yes | Aphids + WW | 1 |
| Chrys_Acet | 94 | 10 | No | No | Control | 2 |
| Chrys_Acet | 94 | 12 | No | No | Control | 9 |
| Chrys_Acet | 94 | 19 | No | No | Control | 1 |
| Chrys_Acet | 94 | 14 | No | Yes | WW | 2 |
| Chrys_Acet | 94 | 16 | No | Yes | WW | 9 |
| Chrys_Acet | 94 | 4 | No | Yes | WW | 1 |
| Chrys_Acet | 95 | 12 | Yes | No | Aphids | 5 |
| Chrys_Acet | 95 | 19 | Yes | No | Aphids | 7 |
| Chrys_Acet | 95 | 2 | Yes | No | Aphids | 6 |
| Chrys_Acet | 95 | 17 | Yes | Yes | Aphids + WW | 7 |
| Chrys_Acet | 95 | 4 | Yes | Yes | Aphids + WW | 6 |
| Chrys_Acet | 95 | 6 | Yes | Yes | Aphids + WW | 5 |

| | | | | | | |
|---|---|---|---|---|---|---|
| Chrys_Acet | 95 | 1 | No | No | Control | 6 |
| Chrys_Acet | 95 | 18 | No | No | Control | 5 |
| Chrys_Acet | 95 | 5 | No | No | Control | 7 |
| Chrys_Acet | 95 | 13 | No | Yes | WW | 5 |
| Chrys_Acet | 95 | 20 | No | Yes | WW | 5 |
| Chrys_Acet | 95 | 3 | No | Yes | WW | 6 |
| Chrys_Acet | 95 | 8 | No | Yes | WW | 7 |
| Chrys_Acet | 100 | 1 | Yes | No | Aphids | 8 |
| Chrys_Acet | 100 | 11 | Yes | No | Aphids | 3 |
| Chrys_Acet | 100 | 16 | Yes | No | Aphids | 4 |
| Chrys_Acet | 100 | 17 | Yes | No | Aphids | 10 |
| Chrys_Acet | 100 | 14 | Yes | Yes | Aphids + WW | 10 |
| Chrys_Acet | 100 | 15 | Yes | Yes | Aphids + WW | 8 |
| Chrys_Acet | 100 | 3 | Yes | Yes | Aphids + WW | 3 |
| Chrys_Acet | 100 | 4 | Yes | Yes | Aphids + WW | 4 |
| Chrys_Acet | 100 | 13 | No | No | Control | 4 |
| Chrys_Acet | 100 | 6 | No | No | Control | 10 |
| Chrys_Acet | 100 | 8 | No | No | Control | 3 |
| Chrys_Acet | 100 | 9 | No | No | Control | 8 |
| Chrys_Acet | 100 | 10 | No | Yes | WW | 3 |
| Chrys_Acet | 100 | 2 | No | Yes | WW | 4 |
| Chrys_Acet | 100 | 20 | No | Yes | WW | 10 |
| Chrys_Acet | 100 | 5 | No | Yes | WW | 8 |
| Athu_Bthu | 55 | 16 | Yes | No | Aphids | 2 |
| Athu_Bthu | 55 | 4 | Yes | No | Aphids | 1 |
| Athu_Bthu | 55 | 5 | Yes | No | Aphids | 9 |
| Athu_Bthu | 55 | 2 | Yes | Yes | Aphids + WW | 2 |
| Athu_Bthu | 55 | 20 | Yes | Yes | Aphids + WW | 9 |
| Athu_Bthu | 55 | 9 | Yes | Yes | Aphids + WW | 1 |
| Athu_Bthu | 55 | 14 | No | No | Control | 1 |
| Athu_Bthu | 55 | 3 | No | No | Control | 2 |
| Athu_Bthu | 55 | 6 | No | No | Control | 9 |
| Athu_Bthu | 55 | 1 | No | Yes | WW | 2 |
| Athu_Bthu | 55 | 11 | No | Yes | WW | 9 |
| Athu_Bthu | 55 | 15 | No | Yes | WW | 1 |
| Athu_Bthu | 56 | 19 | Yes | No | Aphids | 7 |
| Athu_Bthu | 56 | 2 | Yes | No | Aphids | 3 |

| | | | | | | |
|---|---|---|---|---|---|---|
| Athu_Bthu | 56 | 3 | Yes | No | Aphids | 10 |
| Athu_Bthu | 56 | 13 | Yes | Yes | Aphids + WW | 7 |
| Athu_Bthu | 56 | 14 | Yes | Yes | Aphids + WW | 3 |
| Athu_Bthu | 56 | 20 | Yes | Yes | Aphids + WW | 10 |
| Athu_Bthu | 56 | 11 | No | No | Control | 7 |
| Athu_Bthu | 56 | 12 | No | No | Control | 3 |
| Athu_Bthu | 56 | 4 | No | No | Control | 10 |
| Athu_Bthu | 56 | 17 | No | Yes | WW | 7 |
| Athu_Bthu | 56 | 18 | No | Yes | WW | 10 |
| Athu_Bthu | 56 | 9 | No | Yes | WW | 3 |
| Athu_Bthu | 57 | 11 | Yes | No | Aphids | 4 |
| Athu_Bthu | 57 | 13 | Yes | No | Aphids | 5 |
| Athu_Bthu | 57 | 14 | Yes | No | Aphids | 8 |
| Athu_Bthu | 57 | 18 | Yes | No | Aphids | 6 |
| Athu_Bthu | 57 | 15 | Yes | Yes | Aphids + WW | 6 |
| Athu_Bthu | 57 | 16 | Yes | Yes | Aphids + WW | 5 |
| Athu_Bthu | 57 | 5 | Yes | Yes | Aphids + WW | 4 |
| Athu_Bthu | 57 | 7 | Yes | Yes | Aphids + WW | 8 |
| Athu_Bthu | 57 | 1 | No | No | Control | 5 |
| Athu_Bthu | 57 | 2 | No | No | Control | 4 |
| Athu_Bthu | 57 | 6 | No | No | Control | 6 |
| Athu_Bthu | 57 | 9 | No | No | Control | 8 |
| Athu_Bthu | 57 | 10 | No | Yes | WW | 5 |
| Athu_Bthu | 57 | 12 | No | Yes | WW | 4 |
| Athu_Bthu | 57 | 19 | No | Yes | WW | 6 |
| Athu_Bthu | 57 | 3 | No | Yes | WW | 8 |
| Bthu_high | 11 | 4 | Yes | No | Aphids | 1 |
| Bthu_high | 11 | 6 | Yes | No | Aphids | 10 |
| Bthu_high | 11 | 9 | Yes | No | Aphids | 3 |
| Bthu_high | 11 | 15 | Yes | Yes | Aphids + WW | 10 |
| Bthu_high | 11 | 7 | Yes | Yes | Aphids + WW | 3 |
| Bthu_high | 11 | 8 | Yes | Yes | Aphids + WW | 1 |
| Bthu_high | 11 | 1 | No | No | Control | 3 |
| Bthu_high | 11 | 16 | No | No | Control | 10 |
| Bthu_high | 11 | 5 | No | No | Control | 1 |
| Bthu_high | 11 | 10 | No | Yes | WW | 3 |
| Bthu_high | 11 | 12 | No | Yes | WW | 10 |

| | | | | | | |
|---|---|---|---|---|---|---|
| Bthu_high | 11 | 18 | No | Yes | WW | 1 |
| Bthu_high | 14 | 12 | Yes | No | Aphids | 9 |
| Bthu_high | 14 | 13 | Yes | No | Aphids | 6 |
| Bthu_high | 14 | 15 | Yes | No | Aphids | 5 |
| Bthu_high | 14 | 16 | Yes | No | Aphids | 7 |
| Bthu_high | 14 | 17 | Yes | Yes | Aphids + WW | 9 |
| Bthu_high | 14 | 19 | Yes | Yes | Aphids + WW | 6 |
| Bthu_high | 14 | 8 | Yes | Yes | Aphids + WW | 7 |
| Bthu_high | 14 | 9 | Yes | Yes | Aphids + WW | 5 |
| Bthu_high | 14 | 1 | No | No | Control | 9 |
| Bthu_high | 14 | 18 | No | No | Control | 7 |
| Bthu_high | 14 | 4 | No | No | Control | 5 |
| Bthu_high | 14 | 6 | No | No | Control | 6 |
| Bthu_high | 14 | 10 | No | Yes | WW | 7 |
| Bthu_high | 14 | 11 | No | Yes | WW | 9 |
| Bthu_high | 14 | 20 | No | Yes | WW | 6 |
| Bthu_high | 14 | 3 | No | Yes | WW | 5 |
| Bthu_high | 17 | 11 | Yes | No | Aphids | 4 |
| Bthu_high | 17 | 12 | Yes | No | Aphids | 8 |
| Bthu_high | 17 | 13 | Yes | No | Aphids | 2 |
| Bthu_high | 17 | 16 | Yes | Yes | Aphids + WW | 4 |
| Bthu_high | 17 | 2 | Yes | Yes | Aphids + WW | 8 |
| Bthu_high | 17 | 7 | Yes | Yes | Aphids + WW | 2 |
| Bthu_high | 17 | 3 | No | No | Control | 4 |
| Bthu_high | 17 | 4 | No | No | Control | 2 |
| Bthu_high | 17 | 9 | No | No | Control | 8 |
| Bthu_high | 17 | 10 | No | Yes | WW | 4 |
| Bthu_high | 17 | 5 | No | Yes | WW | 2 |
| Bthu_high | 17 | 8 | No | Yes | WW | 8 |
| Mixed_low | 64 | 1 | Yes | No | Aphids | 10 |
| Mixed_low | 64 | 5 | Yes | No | Aphids | 5 |
| Mixed_low | 64 | 8 | Yes | No | Aphids | 4 |
| Mixed_low | 64 | 14 | Yes | Yes | Aphids + WW | 10 |
| Mixed_low | 64 | 18 | Yes | Yes | Aphids + WW | 5 |
| Mixed_low | 64 | 9 | Yes | Yes | Aphids + WW | 4 |
| Mixed_low | 64 | 19 | No | No | Control | 5 |
| Mixed_low | 64 | 2 | No | No | Control | 4 |

| | | | | | | |
|---|---|---|---|---|---|---|
| Mixed_low | 64 | 20 | No | No | Control | 10 |
| Mixed_low | 64 | 3 | No | Yes | WW | 4 |
| Mixed_low | 64 | 4 | No | Yes | WW | 5 |
| Mixed_low | 64 | 6 | No | Yes | WW | 10 |
| Mixed_low | 67 | 1 | Yes | No | Aphids | 8 |
| Mixed_low | 67 | 11 | Yes | No | Aphids | 2 |
| Mixed_low | 67 | 12 | Yes | No | Aphids | 9 |
| Mixed_low | 67 | 18 | Yes | Yes | Aphids + WW | 9 |
| Mixed_low | 67 | 2 | Yes | Yes | Aphids + WW | 2 |
| Mixed_low | 67 | 3 | Yes | Yes | Aphids + WW | 8 |
| Mixed_low | 67 | 10 | No | No | Control | 9 |
| Mixed_low | 67 | 15 | No | No | Control | 8 |
| Mixed_low | 67 | 5 | No | No | Control | 2 |
| Mixed_low | 67 | 16 | No | Yes | WW | 8 |
| Mixed_low | 67 | 20 | No | Yes | WW | 2 |
| Mixed_low | 67 | 7 | No | Yes | WW | 9 |
| Mixed_low | 68 | 10 | Yes | No | Aphids | 1 |
| Mixed_low | 68 | 12 | Yes | No | Aphids | 3 |
| Mixed_low | 68 | 14 | Yes | No | Aphids | 6 |
| Mixed_low | 68 | 18 | Yes | No | Aphids | 7 |
| Mixed_low | 68 | 13 | Yes | Yes | Aphids + WW | 6 |
| Mixed_low | 68 | 15 | Yes | Yes | Aphids + WW | 3 |
| Mixed_low | 68 | 17 | Yes | Yes | Aphids + WW | 1 |
| Mixed_low | 68 | 9 | Yes | Yes | Aphids + WW | 7 |
| Mixed_low | 68 | 1 | No | No | Control | 6 |
| Mixed_low | 68 | 11 | No | No | Control | 3 |
| Mixed_low | 68 | 16 | No | No | Control | 7 |
| Mixed_low | 68 | 8 | No | No | Control | 1 |
| Mixed_low | 68 | 2 | No | Yes | WW | 7 |
| Mixed_low | 68 | 3 | No | Yes | WW | 6 |
| Mixed_low | 68 | 6 | No | Yes | WW | 3 |

**Table S6:** R-code for all models

| Table | Model | R-code |
|---|---|---|
| 1 | A | Lmer(sqrt(Aphid colony size) ~ Belowground |

| | | |
|---|---|---|
| | | (yes/no) * Chemotype * Day<br>+ (1\|Block) + (1\|Daughter) + (1\|Day/Daughter) |
| 1 | B | Lmer(sqrt(Aphid colony size) ~ Retrieved wireworm larvae * Chemotype * Day<br>+ (1\|Block) + (1\|Daughter) + (1\|Day/Daughter) |
| 2 | A | Lmer(sqrt(Final aphid colony size) ~ Belowground (yes/no) + Terpenoid evenness + Terpenoid richness + Terpenoid concentration<br>+ (1\|Block) |
| 2 | B | Lmer(sqrt(Final aphid colony size) ~ Retrieved wireworm larvae + Terpenoid evenness + Terpenoid richness + Terpenoid concentration<br>+ (1\|Block) |
| 3 | Plant dry weight | Lmer(Dry weight ~ *Coloradoa tanacetina* + Aboveground * Belowground * Chemotype<br>+ (1\|Block) + (1\|Daughter) |
| 3 | Plant height | Lmer(Plant height ~ *Coloradoa tanacetina* + Aboveground * Belowground * Chemotype<br>+ (1\|Block) + (1\|Daughter) |
| 3 | Chlorophyll content | Lmer(Chlorophyll content ~ *Coloradoa tanacetina* + Aboveground * Belowground * Chemotype<br>+ (1\|Block) + (1\|Daughter) |
| 7 | Chemotype | Glmer(Aphid survival ~ Chemotype<br>+ (1\|Block) + (1\|Daughter),<br>Family = binomial |
| 7 | Daughters | Glmer(Aphid survival ~ (1\|Chemotype/Daughter)<br>+ (1\|Block), |

| | | |
|---|---|---|
| | | Family = binomial |
| 8 | A, B | See Table 1; Plants with zero aphids were included into the dataset |
| 9 | A, B | See Table 2; Plants with zero aphids were included into the dataset |
| 10 | A | Lmer(sqrt(Final aphid colony size) ~ *Coloradoa tanacetina* + Chemotype * Belowground (yes/no) + (1\|Block) + (1\|Daughter) |
| 10 | B | Lmer(sqrt(Final aphid colony size) ~ *Coloradoa tanacetina* + Chemotype * Retrieved wireworm larvae + (1\|Block) + (1\|Daughter) |
| 11 | | Lmer(sqrt(*Coloradoa tanacetina*) ~ Chemotype * sqrt(Final aphid colony size) + (1\|Block) + (1\|Daughter) |

**Table S7:** Output from a mixed linear model for *M. tanacetaria* survival in plants that were assigned aphid treatment, using chemotype (Model A) or daughter nested within chemotype (Model B) as fixed effect and the daughter (only in Model A) and the block as random effects.

| | | **d.f.** | **$\chi^2$ (p-value)** |
|---|---|---|---|
| **Model A** | Chemotype | 5 | 7.36 (0.195) |
| **Model B** | Daughter | 1 | **11.30 (<0.001)** |

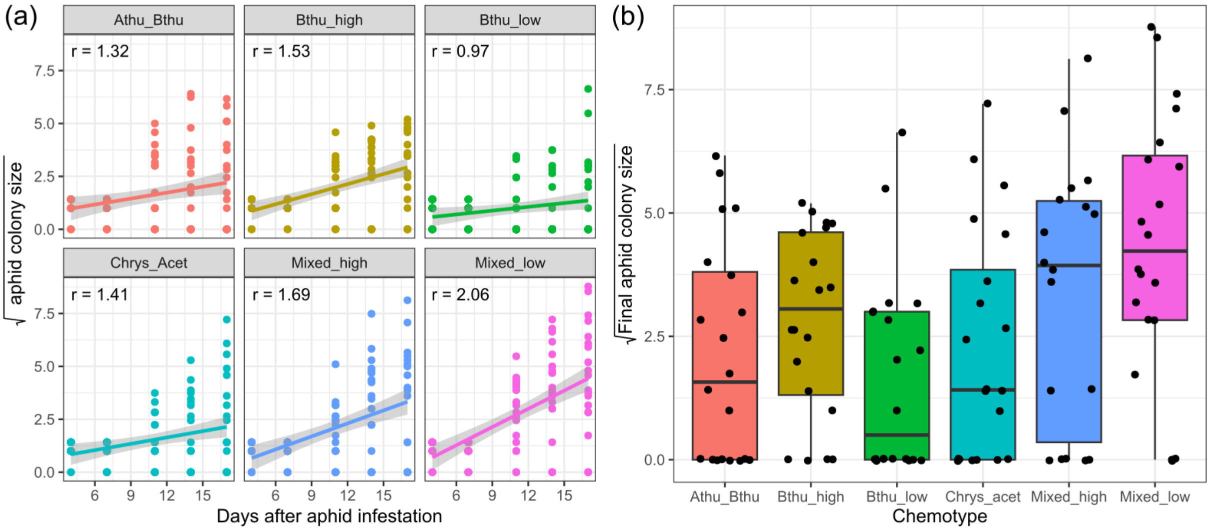


**Fig. S5** (a) Square root-transformed *M. tanacetaria* colony size over time in days after aphid infestation, across chemotypes. R-values represent the slope of fitted lines. (b) Final aphid colony size at the time of the experimental harvest for different tansy chemotypes. Boxes represent the variation in data, where the lower hinge corresponds to the first quartile (25th percentile) and the upper hinge depicts the third quartile (75th percentile). Whiskers indicate the 5% and 95% percentiles; solid lines within boxes represent the medians. Black dots indicate individual sample values. The six chemotypes are depicted in different colours for convenience

**Table S8:** Output from a mixed linear model for *M. tanacetaria* colony size over time, using either belowground herbivory treatment (Belowground; Model A) or the number of retrieved wireworm larvae (Wireworm larvae; Model B), and day and chemotype, and the interaction terms as fixed effects. In both models, block, daughter ID, and individual ID (nested within day) were used as random effects.

| **Model A** | **d.f.** | **$\chi^2$ (p-value)** | **Model B** | **d.f.** | **$\chi^2$ (p-value)** |
|---|---|---|---|---|---|
| Belowground | 1 | 2.04 (0.153) | Wireworm larvae | 3 | 2.95 (0.399) |
| Chemotype | 5 | 7.33 (0.197) | Chemotype | 5 | 6.67 (0.246) |
| Day | 1 | **86.01 (<0.001)** | Day | 1 | **84.71 (<0.001)** |
| Belowground * Chemotype | 5 | 3.06 (0.691) | Wireworm larvae * Chemotype | 14 | 13.30 (0.503) |
| Belowground * | 1 | 0.10 (0.751) | Wireworm larvae | 3 | 4.03 (0.258) |

| | | | | | |
|---|---|---|---|---|---|
| Day | | | * Day | | |
| Chemotype * Day | 5 | **22.85 (0.001)** | Chemotype * Day | 5 | **17.34 (0.004)** |
| Belowground * Chemotype * Day | 5 | 3.62 (0.606) | Wireworm larvae * Chemotype * Day | 14 | 9.02 (0.830) |

**Table S9:** Output from a mixed linear model for final *M. tanacetaria* colony size, using belowground herbivory treatment (BG; Model A) or the number of retrieved wireworm larvae (Wireworm larvae; Model B) and terpenoid richness, terpenoid evenness and total terpenoid concentration calculated based on the terpenoid profile of the 18 daughter plants (three for each of the six chemotypes) as fixed effects and the block as random effect.

| **Model A** | **d.f.** | **$\chi^2$ (p-value)** | **Model B** | **d.f.** | **$\chi^2$ (p-value)** |
|---|---|---|---|---|---|
| BG | 1 | 0.52 (0.469) | Wireworm larvae | 3 | **8.56 (0.036)** |
| Evenness | 1 | **5.77 (0.016)** | Evenness | 1 | *3.82 (0.051)* |
| Richness | 1 | 0.00 (0.990) | Richness | 1 | 0.00 (0.951) |
| Concentration | 1 | 2.00 (0.157) | Concentration | 1 | 2.18 (0.140) |

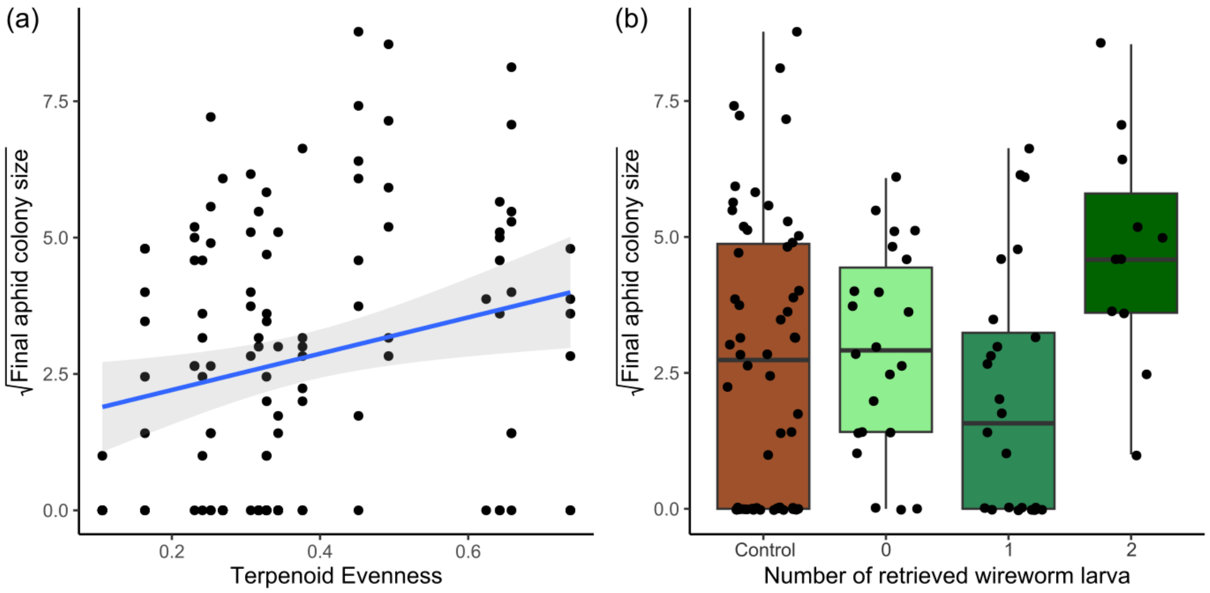


**Fig. S6** (a) Square root-transformed *M. tanacetaria* colony size on plants differing in leaf terpenoid evenness. The linear trendline depicts average predicted values based on a linear model, and the shaded area depicts the 95% confidence interval. (b) Box plots visualizing square root-transformed *M. tanacetaria* colony size on plants with no added wireworms, compared to plants on which 0, 1 or 2 wireworm larvae were retrieved after the harvest. Boxes represent the variation in data, where the lower hinge corresponds to the first quartile (25th percentile) and the upper hinge depicts the third quartile (75th percentile). Whiskers indicate the 5% and 95% percentiles; solid lines within boxes represent the medians. Black dots indicate individual sample values

**Table S10:** Output from a mixed linear model for the *M. tanacetaria* colony size (square root transformed), taking *C. tanacetina* abundance, chemotype, and either belowground herbivory treatment (BG; Model A) or the number of retrieved wireworm larvae (Wireworm larvae; Model B) and the interplay of chemotype and belowground treatment or chemotype and the number of retrieved wireworm larvae as fixed effects, and the daughter and block as random effect into account.

| **Model A** | **d.f.** | **$\chi^2$ (p-value)** | **Model B** | **d.f.** | **$\chi^2$ (p-value)** |
|---|---|---|---|---|---|

| | | | | | |
|---|---|---|---|---|---|
| *C. tanacetina* | 1 | 2.09 (0.148) | *C. tanacetina* | 1 | 1.53 (0.216) |
| Chemotype | 5 | **14.66 (0.012)** | Chemotype | *5* | *9.40 (0.094)* |
| BG | 1 | 0.17 (0.683) | Wireworm larvae | 3 | 5.79 (0.122) |
| Chemotype * BG | 5 | 3.28 (0.657) | Chemotype * Wireworm larvae | 14 | 9.45 (0.801) |

**Table S11:** Output from a mixed linear model for the *C. tanacetina* abundance (Square root transformed), taking *M. tanacetaria* colony size (square root transformed), chemotype, and the interplay of chemotypes and *M. tanacetaria* as fixed effects, and the daughter and block as random effect into account.

| | **d.f.** | **$\chi^2$ (p-value)** |
|---|---|---|
| Chemotype | 5 | 1.49 (0.914) |
| *M. tanacetaria* | 1 | 2.21 (0.137) |
| Chemotype * *M. tanacetaria* | 5 | 7.66 (0.176) |

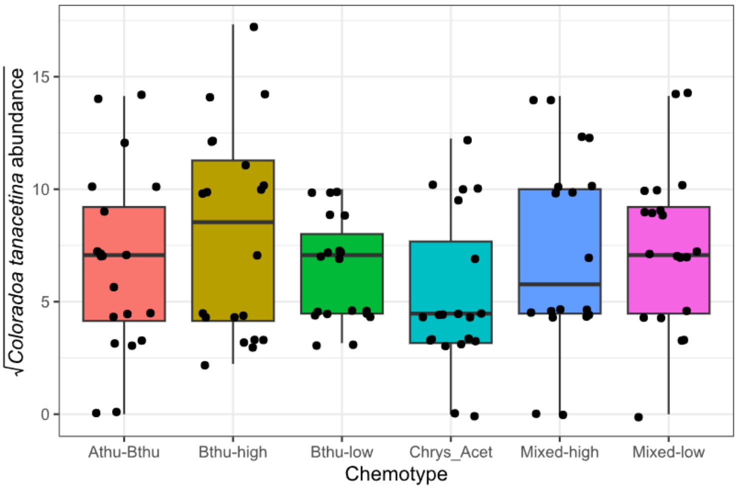


**Fig. S7** Estimated *C. tanacetina* numbers for different tansy chemotypes three days after the final count of *M. tanacetaria*. Boxes represent the variation in data, where the lower hinge corresponds to the first quartile (25th percentile) and the upper hinge depicts the third quartile (75th percentile). Whiskers indicate the 5% and 95% percentiles; solid lines within boxes represent the medians. Black dots indicate individual sample values. The six chemotypes are depicted in different colours for convenience

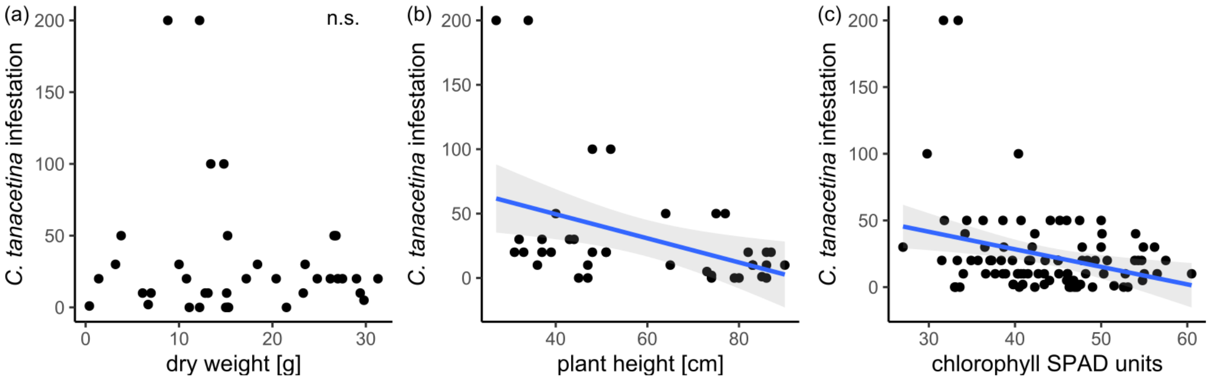


**Fig. S8** (a) Infestation of *Coloradoa tanacetina* in relationship to plant dry weight (g). (b) Infestation of *Coloradoa tanacetina* in relationship to plant height (cm). (c) Infestation of *Coloradoa tanacetina* in relationship to plant chlorophyll (SPAD units). The linear trendline in (b) and (c) depicts average predicted values based on a linear model, and the shaded area depicts the 95% confidence interval
